# Correlation of gene expressions between nucleus and cytoplasm reflects single-cell physiology

**DOI:** 10.1101/206672

**Authors:** Mahmoud N. Abdelmoez, Kei Iida, Yusuke Oguchi, Hidekazu Nishikii, Ryuji Yokokawa, Hidetoshi Kotera, Sotaro Uemura, Juan G. Santiago, Hirofumi Shintaku

## Abstract

**Background:** Eukaryotes transcribe RNAs in nuclei and transport them to the cytoplasm through multiple steps of post-transcriptional regulation. Existing single-cell sequencing technologies, however, are unable to analyse nuclear (nuc) and cytoplasmic (cyt) RNAs separately and simultaneously. Hence, there remain challenges to discern correlation, localisation, and translocation between them.

**Results:** Here we report a microfluidic system that physically separates nucRNA and cytRNA from a single cell and enables single-cell integrated nucRNA and cytRNA-sequencing (SINC-seq). SINC-seq constructs two individual RNA-seq libraries, nucRNA and cytRNA per cell, quantifies gene expression in the subcellular compartments and combines them to create a novel single-cell RNA-seq data enabled by our system, which we here term in-silico single cell.

**Conclusions:** Leveraging SINC-seq, we discovered three distinct natures of correlation among cytRNA and nucRNA that reflected the physiological state of single cells: The cell-cycle-related genes displayed highly correlated expression pattern with minor differences; RNA splicing genes showed lower nucRNA-to-cytRNA correlation, suggesting a retained intron may be implicated in inhibited mRNA transport; A chemical perturbation, sodium butyrate treatment, transiently distorted the correlation along differentiating human leukemic cells to erythroid cells. These data uniquely provide insights into the regulatory network of mRNA from nucleus toward cytoplasm at the single cell level.

## Background

Single-cell sequencing is a powerful tool to explore epigenetic, genomic and transcriptional heterogeneities at unprecedented resolution[1–6]. RNA-seq with single nuclei (nucRNA-seq) is an emerging alternative to profile gene expressions of cells in tissues that cannot be readily dissociated such as the adult brain and frozen samples. The method is further capable of coupling with sorting by fluorescence activated cell sorters [4, 7], Fluidigm C1[5], and Drop-seq[8], and demonstrated feasibilities of identifying cell types and cell cycles with nucRNA-seq data[9]. Although these works hypothesise that the nucRNA expression is representative of whole cells, to date, the direct evidence of the correlation in the cytRNA and nucRNA expression at single-cell resolution has not been provided.

Recent technical advances have further enabled combined sequencing at multi-omic levels within the same single cells[10–12] and helped to understand the links underlying the regulatory cascade. Several microfluidic[13–16] and non-microfluidic protocols[17, 18] offer parallel transcriptional and genomic analyses on the same single cell by fractionating cytRNAs and nuclei of single cells. However, we know of no work that has reported an integrated nucRNA-seq and cytRNA-seq with the same cell to study RNA transport and gene regulation and function through splicing of pre-mRNA[19, 20].

Here we develop a novel single-cell sequencing method, SINC-seq, which combines a microfluidic protocol that physically fractionates nuclear and cytoplasmic RNAs and a subcellular RNA-seq pipeline to dissect RNA expressions in the individual subcellular compartment. We utilise SINC-seq to explore both correlated and uncorrelated gene expression between the compartments with single K562 human leukemic cells. We further explore correlation dynamics that reflect the transient response of cells with differentiating K562 cells to erythroid cells under a perturbation of sodium butyrate, a histone deacetylase inhibitor. These data reveal how eukaryotes manage subcellular RNA expressions via inter-compartment regulation.

## Results

### A Microfluidic platform for single-cell integrated nuclear and cytoplasmic RNA-sequencing: SINC-seq

To dissect transcriptional correlation in the subcellular compartments, we devised SINC-seq that combines an electrophoretic fractionation of cytRNA from the nucleus[14–16] with off-chip RNA sequencing (Fig. 1a, b). SINC-seq constructs individual RNA-seq libraries with cytRNA and nucRNA and integrates the sequencing data in a new form of sequencing data, which we term an in-silico single cell. SINC-seq starts with a microfluidic protocol that leverages a hydrodynamic trap that captures a single cell, concentrates an electrical field to lyse the cytoplasmic membrane selectively while leaving the nuclear membrane relatively intact, and retains the nucleus during electric-field-based extraction of cytRNA to fractionate them (Fig.1b, c, Supplementary Fig. 1, Supplementary Video 1, and Methods). The microfluidic system completes the entire process with a voltage control via three end-channel electrodes and outputs the cytRNA and nucleus to different wells in less than 5 min. We note that the hydrodynamic trap integrated in this work couples hydrodynamic flow and electric field concentration, and uniquely enables a highly automated workflow and about 20-fold reduction in the applied voltage compared to our previous protocol[14–16]. These key improvements allowed us to study RNA expressions in subcellular compartments of single cells systematically.

**Figure 1.**
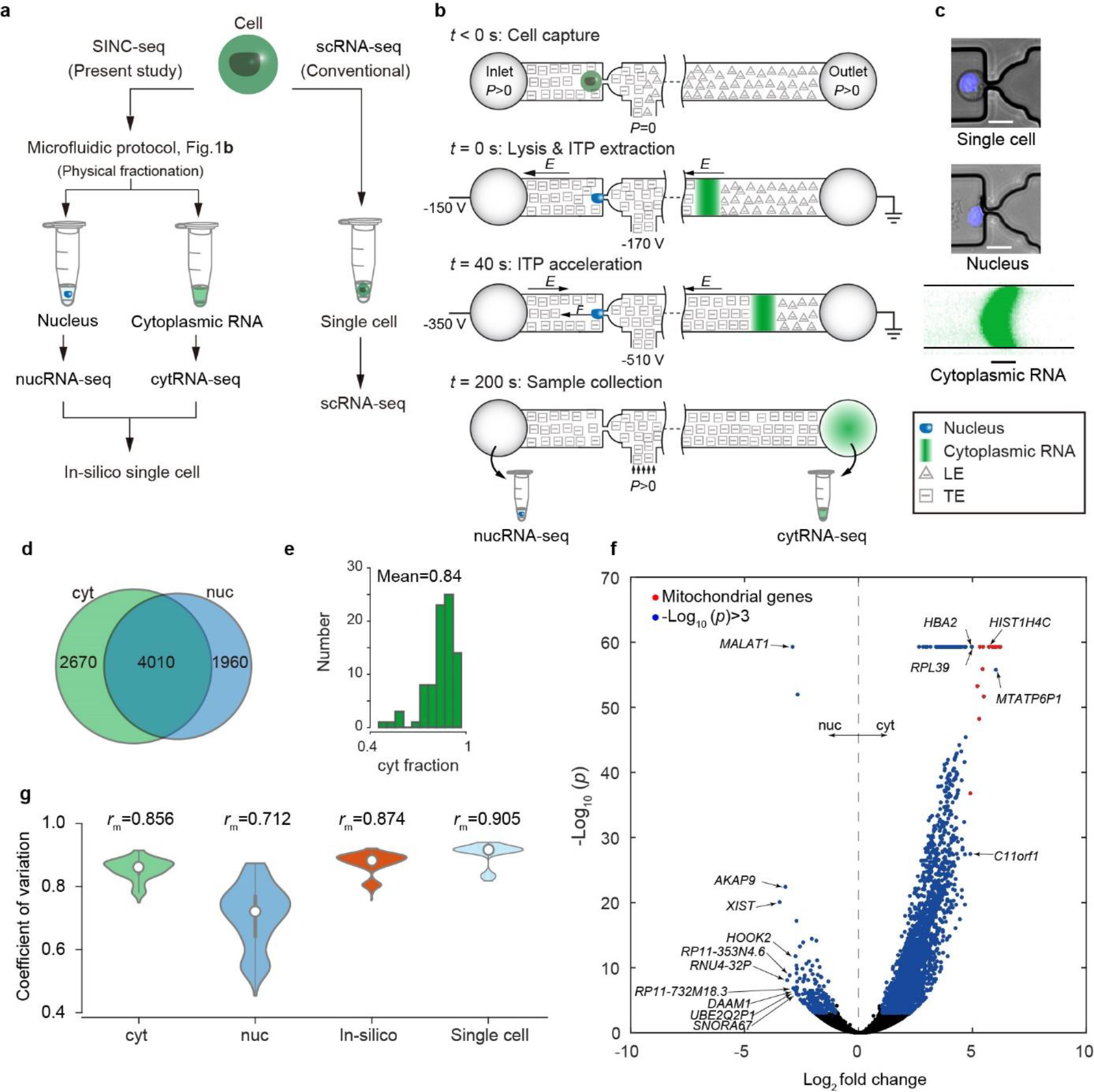
Single-cell integrated nuclear and cytoplasmic RNA-seq (SINC-seq). a, SINC-seq and conventional scRNA-seq. b, Workflow of SINC-seq. Single cell isolation at a hydrodynamic trap via pressure-driven flow (*t*=0 s); Lysis of cell membrane and cytRNA extraction with isotachophoresis (ITP)-aided nucleic acids extraction (*t*>0 s); ITP acceleration by changing voltages (*t*=40 s); Voltage deactivation and sample collection from the wells of the microchannel (*t*>200 s). c, Fluorescence microscopy images of the trapped single-cell, nucleus after cytRNA extraction (stained with Hoechst) and extracted cytRNA stained with SYBR Green II. The bars are 20 μm. d, Venn diagram of mean numbers of detected genes in cytRNA-seq and nucRNA-seq. e, Proportion of abundance of transcripts in the cytoplasm. f, Differential expression between cytRNA and nucRNA. Blue, genes with *p* values less than 0.001 and absolute log2 fold changes greater than unity. g, Correlation coefficients of gene expression pattern computed with respect to the conventional scRNA-seq, showing our novel in-silico single cell normalisation showed the best correlation with the scRNA-seq.

### Library preparation and quality control with SINC-seq

To critically evaluate SINC-seq, we performed experiments with 93 single cells of K562 human myeloid leukaemia cells and generated 186 RNA-seq libraries with off-chip Smart-seq2 protocol[21]. Of the 93 single cells analysed in this experiment, all showed successful extraction with monitoring current during extraction (Supplementary Fig. 1c); 84 (QC) for both cytRNA-seq and nucRNA-seq. Of the 93 single cells analyzed in this experiment, 9 failed quality control (QC) for either cytRNA-seq or nucRNA-seq; in 7 of the samples that failed QC, we observed low yield in the amplification either with cytRNA or nucRNA; in two of the samples, we observed incomplete fractionation (see Supplementary Figs. 2-4 and Supplementary Data 1). After the QC, we achieved 168 data sets consisting of 84 pairs of cytRNA-seq and nucRNA-seq. Our protocol showed lower cDNA yield with a nucleus than with cytRNA and the total cDNA amount of the cDNA per single cell was 26±16% less than that with a conventional single-cell protocol on average (Supplementary Fig.2a), however this does not result in a significant decrease in the sensitivity and repeatability (see Supplementary Information and Supplementary Fig.3).

### SINC-seq dissects the difference in subcellular gene expression

To assess the performance of SINC-seq, we computed gene expression with an in-silico single cell analysis. We used this to benchmark the sensitivity and repeatability of our methods (Methods, see a comprehensive benchmark of SINC-seq in Supplementary Figs. 3 and 4 and the Supplementary Information). SINC-seq consistently detected 6,680±1,360 (mean±s.d.) and 5,970±1,400 genes per cytRNA and nucRNA, respectively, and 8,640±1,150 genes per cell with transcripts per million (TPM) greater than 0 (Fig. 1d). SINC-seq also revealed that ~16% transcripts were in the nucleus and ~84% in the cytoplasm (Fig. 1e) and enriched expression of 226 and 3,035 genes, respectively (Fig. 1f). On average, SINC-seq displayed about 6.3% smaller number of detected genes than the conventional single-cell RNA-seq (scRNA-seq) that detected 9,230±1,220 genes (n=12). Notably, our in-silico single cell data showed a wider dynamic range in the detection of genes as compared to scRNA-seq (Supplementary Fig. 4k). To the scRNA-seq, the in-silico single-cell data showed the higher coefficient of correlation computed with log-transformed expression (log10(TPM+1)) than either cytRNA-seq or nucRNA-seq (Fig. 1g). Combined, average gene expression profile of 12 in-silico single-cells (Supplementary Fig. 5) showed an excellent matching with that of 12 scRNA-seq (*r* = 0.949, Pearson correlation coefficient, Supplementary Fig. 3q). The total number of detected genes in the 12 in-silico single cells was 15,314, of which 13,286 genes were also detected in the average of the 12 scRNA-seq (Supplementary Fig. 3t). We again stress that the in-silico normalisation and resulting scRNA-seq is a novel method, which uniquely leverages the physical separation and minimal cross-contamination enabled by our electrophoretic fractionation.

### Cell cycle-related genes show correlated expression in cytoplasm and nucleus

To view the landscape of the correlation between nucRNA and cytRNA, we computed cross-correlation of the individual gene as a measure of covariation in the two subcellular compartments and ranked the genes with the coefficient of correlation (Fig. 2a). Gene ontology analysis revealed that the highly correlated genes (top 10%) had cell-cycle as an enriched function (Fig. 2b) and the bottom 10% including anti-correlation had RNA processing (Fig. 2c).

**Figure 2.**
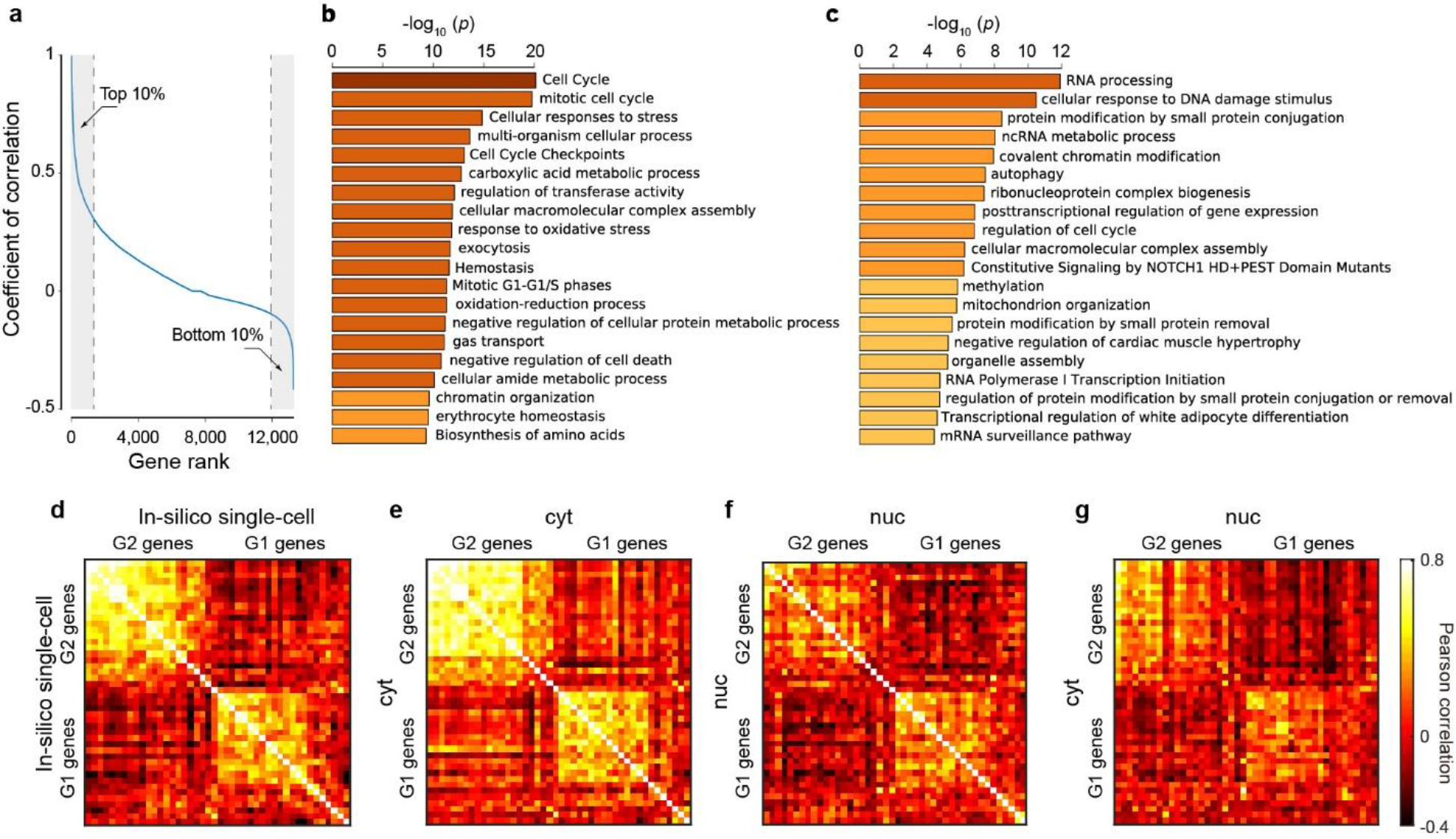
Landscape of cross-correlation between cytRNA and nucRNA unveiled transcriptional oscillation of cell-cycle genes in nucRNA highly correlated with expression in cytRNA. a, Quantile plot of genes with coefficients of cross-correlation, b Gene ontology analysis with top 10% genes in the quantile plot and c, bottom 10% genes. d-f, Cell-cycle genes in in-silico single cell data, cytRNA, and nucRNA show correlation with in-phase genes (G1 vs. G1 or G2 vs. G2) and anti-correlation with out-of-phase genes (G1 vs. G2). g, The transcriptional oscillation of cell-cycle genes in nucRNA cross-correlated with the gene expression in cytRNA.

To dissect the highly correlated gene expression, we first focused on cell cycle based on transcriptional oscillations[2] and phase-score analysis[3] (Supplementary Information). The in-silico single cell data showed the progression of the cell cycle with a correlated variation of in-phase genes and anti-correlation out-of-phase genes (G1 versus G2) (Fig. 2d), consistent with scRNA-seq of K562[2], and also with the progression of the phase-score (Supplementary Fig. 6a). Similarly, both of cytRNA-seq and nucRNA-seq data revealed the cell cycle (Fig. 2e, f and Supplementary Fig. 6b, c). Notably, we found that the anti-correlation among out-of-phase genes was slightly higher in a nucRNA (U-test, *p* = 7.81×10^−26^, Supplementary Fig. 6d), suggesting that nucRNA-seq more directly detected phase transition in the transcriptional oscillation of cell-cycle genes. On the other hand, the correlation among in-phase genes was higher in cytRNA may indicate further modulation in the cytoplasm.

To elucidate how the transcriptional oscillations in nucRNA modulated the gene expression in the cytRNA, we extended the analyses to compute the cross-correlation between cytRNA and nucRNA with cell-cycle genes. The cell-cycle genes showed synchronised oscillation in the each of the two subcellular compartments (Fig. 2g), consistent with the observation that the subpopulations segregated into the G1 and G2 groups showed corresponding up- and down-regulation of G1 and G2 genes (Supplementary Fig. 6e,f, Supplementary Information). Together, these results suggested that the cytRNA and nucRNA had similar expression patterns of cell-cycle genes and both of them solely had a potency to detect the cell-cycle.

### Nuclear retained intron attenuates the transcriptional oscillation

As an instance of the uncorrelated genes, we next studied the retained intron (RI) mediated regulation of mRNA transport[22–24] leveraging the intron-rich reads with nucRNA-seq of SINC-seq (see Supplementary Fig. 7 for comprehensive statistics on intron detection). After filtering RI (Methods), SINC-seq detected 2,000±740 RIs per cell, of which 1,480±698 and 814±330 were detected with nucRNA-seq and cytRNA-seq, respectively (Fig. 3 a). We identified 223 nuclear-retained introns (NRI) in 202 genes (Methods). Gene ontology analysis[25] found that the 202 genes had enriched functions like RNA processing and RNA splicing (Fig. 3b), consistent with the previous studies[23, 26, 27]. We examined the relationship between the probability of NRI and gene expression in an individual fraction. We observed a positive correlation between them in the nucRNA (*r*=0.436, *p*<0.01 Fig. 3c), while no correlation in the cytRNA (*r*=0.063, *p*=0.439, Fig. 3d). In contrast, we observed different enriched functions with cytoplasmic enriched RIs (CRI) and no correlation between probabilities of CRI and gene expressions (Supplementary Fig. 8a-c). For a better understanding of the function of NRI, we examined the expression patterns of top seven genes that were highly associated with NRI mediated regulation (Fig. 3e-g and Supplementary Figs. 8d-h). Notably, the seven genes contained three splicing-related genes[28] and two snoRNA host genes[29]. These data lead us to hypothesise the NRIs likely attenuates the transcriptional oscillation in the nucleus via fine-tuning RNA metabolism in order to maintain the gene expression and functional RNAs in the cytoplasm.

**Figure 3.**
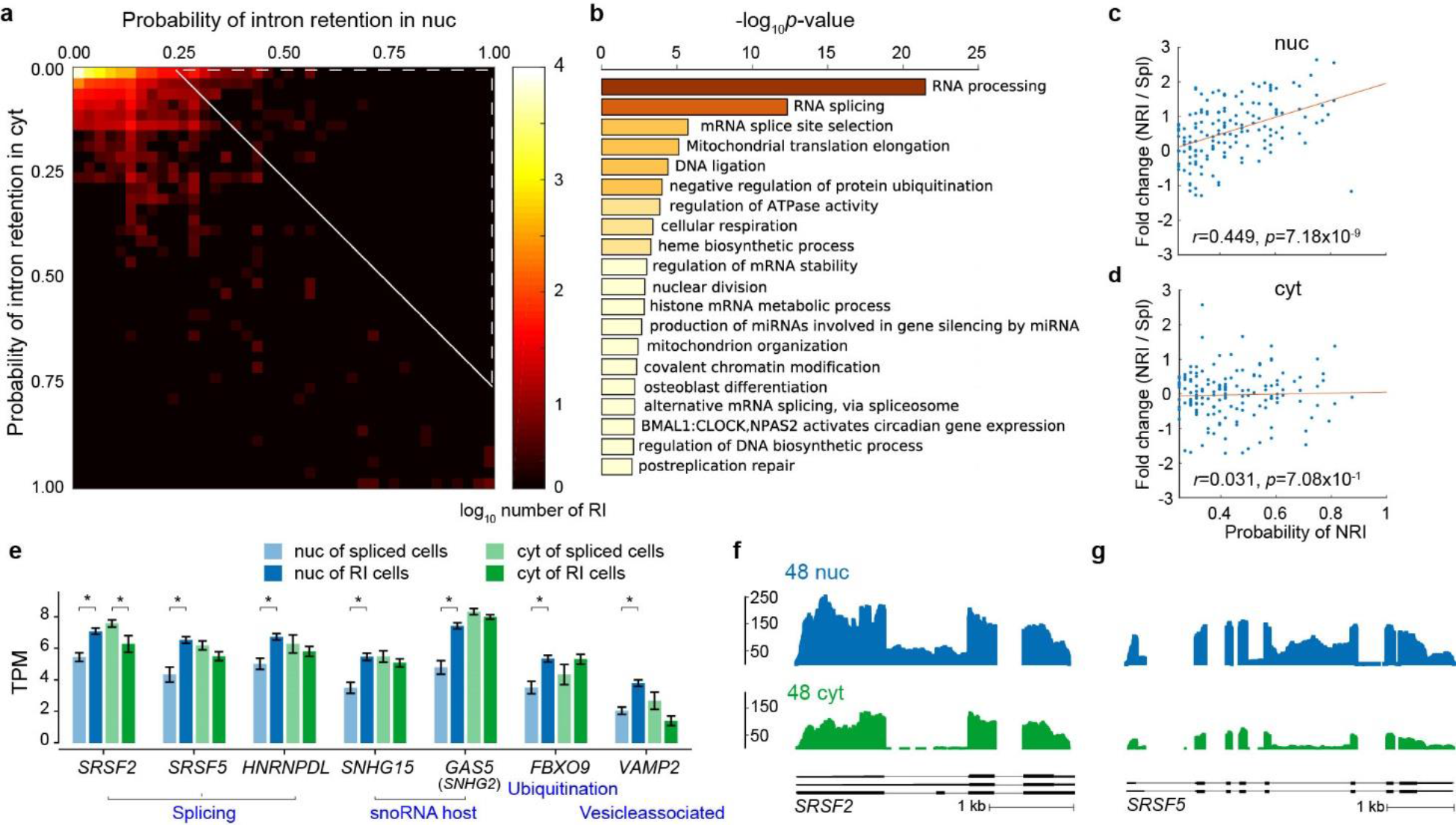
NRI mediated attenuation of transcriptional oscillation in nucRNA. a, Heatmap of RI with the probability of RI in cytRNA and nucRNA fractions. NRI was identified in the upper right region indicated with the broken white line. b, Gene ontology analysis with NRI. c, d, Correlation analysis between the probability of NRI and the fold change of gene expression among cells with NRI and without NRI (Spl: spliced) in nucRNA and cytRNA, respectively. e, Expression of top seven genes that were highly regulated by NRI in nucRNA (*p* <0.001, Mann-Whitney U test), comparing with NRI versus without NRI (Spl) in an individual fraction. f, Coverage of *SRSF2* and g, *SRSF5* genes showing higher intron reads in the nuclear fraction. Coverages of *HNRNPDL*, *SNHG15*, *GAS5* (*SNHG2*), *FBXO9*, and *VAMP2* genes are provided in Supplementary Fig. 8.

### Sodium butyrate treatment on K562 drives diverging gene expression in subcellular compartments

To explore the correlation dynamics under perturbation, we further performed SINC-seq with differentiating K562 cells to erythroid cells by sodium butyrate (Methods), sampling 8-13 cells per day over five days. With 41 successful SINC-seq datasets (82 RNA-seq data in total), we detected differentially expressed genes (DEG), 264 up- and 177 down-regulated genes were in cytRNA, and 64 up-and two down-regulated genes were in nucRNA (Supplementary Fig. 9). We examined the dynamics of the cross-correlation of DEG expression between cytRNA and nucRNA along with the pseudo-time[30](Fig. 4a), which was obtained with in-silico single cell data (Supplementary Fig. 10a). The cross-correlation between cytRNA and nucRNA within the same single cell (along with the diagonal shown in Fig. 4a) exhibited gradual decrease with the pseudo-time, suggesting that the subcellular gene expression patterns behaved differently and diverged along the differentiation. Notably, the correlation between nucRNA fourth day and cytRNA first day (near the top right corner in Fig. 4a) displayed lower correlation as compared to that between nucRNA first day and cytRNA fourth day (near the bottom left corner), suggesting strongly that nucRNA was the driver of the diverging gene expression. We reinforced this observation with the control experiments (Supplementary Fig. 10b-d).

**Figure 4.**
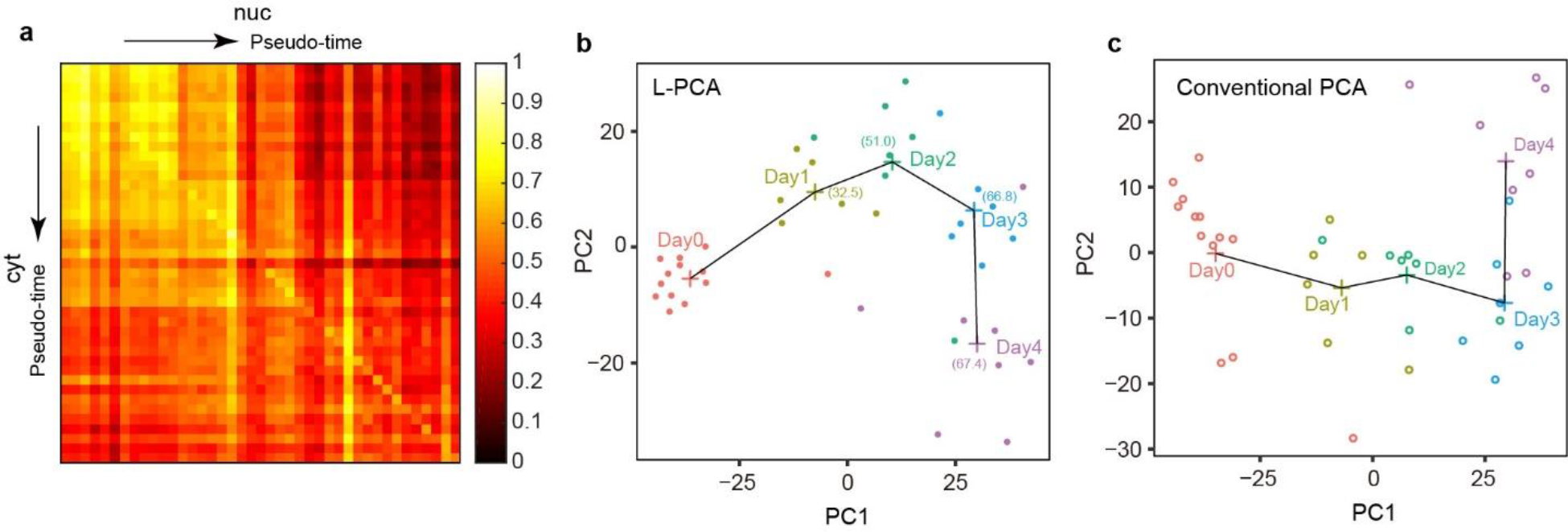
Differentiation of K562 cells to erythroid cells shows a dynamical change of crosscorrelation between cytRNA and nucRNA. a, The cross-correlation of DEG expression between cytRNA and nucRNA along with the pseudo-time. b, L-PCA analysis of differentiating K562 cells to erythroid cells by sodium butyrate treatment. c, Conventional PCA analysis.

We hypothesised that the divergence of gene expressions in the two subcellular compartments might reflect the transient response of different regulatory pathways and enable to resolve the on/off regulation of the transcription leveraging nucRNA-seq. To test this hypothesis, we introduced a localisation-embedded principal components analysis (L-PCA) that computed PCs with the subcellular gene expression of DEGs (Fig. 4b). As expected, L-PCA resolved the trajectory of the differentiation slightly clearer than a conventional PCA that computed PCs with in-silico single cell data (Fig. 4c). To further corroborate the L-PCA, we performed PCA on datasets of an individual fraction, showing this specific and unique transient in the nucRNA compared to that in the cytRNA— that is, in nucRNA, the third day cluster was furthest from the day 0^th^ cluster. (Supplementary Fig. 10e, f). These results demonstrated that the SINC-seq was able to detect important regulatory events. Such events are usually masked by abundant cytoplasmic transcripts in conventional single-cell sequencing.

## Discussion

A fundamental question is how the transcriptional oscillation in the nucRNA, which is inherently stochastic, is transported to and correlated with gene expression in the cytRNA. SINC-seq enabled direct and quantitative comparison of gene expressions between a nucleus, a cytoplasm and a whole cell of a same single cell, revealing that the cells may conceivably fine-tune a portion of their expression upon transport to the cytoplasm (e.g. NRI genes), while preserving correlation of other portions of their expression upon transport (e.g. cell cycle-related genes). SINC-seq also revealed that the cells under the external perturbation dynamically alter the correlation and exhibit the unique trajectory of differentiation at subcellular resolution. These findings shed new light on the characteristics of post-transcriptional regulations with a single cell and subcellular compartment resolution.

Our study also suggests a compelling caution to an approach that approximates the transcriptomic profile of the whole cell with that of a single compartment without validation. The SINC-seq platform will be broadly applicable to different types of cells as long as they are isolated as singles. The method thus will contribute to validate existing subcellular RNA-seq methods[4, 5, 7–9, 17, 18] and also define their limitations.

## Conclusion

To dissect transcriptional correlation in the subcellular compartments, we devised SINC-seq that enables integrated nuclear and cytoplasmic RNA-seq of single cells by coupling a physical fractionation of cytRNA from the nucleus of a single cell with a high-throughput RNA-seq. Leveraging SINC-seq, we explored the landscape of the correlation between nucRNA and cytRNA with a total of 84 K562 cells, which corresponds to 168 RNA-seq libraries. The SINC-seq data unveiled three distinct natures of correlation among cytRNA and nucRNA that reflected the physiological state of single cells: highly-correlated expression in cell-cycle-related genes, the distorted correlation via nuclear-retained intron, and the correlation dynamics along the differentiation of K562 cells to erythroid cells under sodium butyrate perturbation. These data uniquely provide insights into the regulatory network of mRNA from nucleus toward cytoplasm at the single cell level.

## Methods

### Cell

We purchased K562 cells (human lymphoblast, chronic myelogenous leukaemia) from RIKEN BioResource Center and JCRB cell bank. We cultured the K562 cells in RPMI-1640 Medium (Life Technologies) with 10% fetal bovine serum and 1% penicillin-streptomycin-glutamine at 37°C in 5% CO_2_. We washed the cells with phosphate buffered saline once and suspended in a sample buffer containing 50 mM imidazole, 25 mM HEPES, and 175 mM sucrose (pH 7.6) at the concentration of ~5 cells/μL and stored on ice until the experiments were performed. To differentiate K562 cells, we incubated K562 cells with 1 mM NaB (sodium butyrate, Sigma-Aldrich, B5887) and harvested after 96 h of induction.

### Buffers

We designed buffers for isotachophoresis (ITP)-based selective extraction, separation (from the trapped nucleus), purification, and transport of cytRNA to the cytRNA output well of the chip (see more detail in Shintaku et al.^13^ and Kuriyama et al.^14^). The leading electrolytes (LE) were 50 mM Tris and 25 mM HCl containing 0.4% a poly(vinylpyrrolidone) (PVP) (calculated pH of 8.1). The trailing electrolytes (TE) were 50 mM Imidazole and 25 mM HEPES containing (initial calculated pH of 8.3) 0.4% PVP. We included PVP to suppress electroosmotic flow. We purchased Tris, HEPES, Imidazole, and HCl from Sigma-Aldrich, and PVP (MW 1 MDa) from Polyscience. We prepared all solutions in UltraPure DNase-/RNase-free deionised (DI) water (Life Technologies).

### Microfluidic system setup

We fabricated polydimethylsiloxane (PDMS, Sylgard 184, Dow Corning) microchannel superstructures (Supplementary Fig. 1a,b) with a soft-lithography and bonded on a glass substrate. SU-8 (SU-8 2025, MicroChem) moulds were prepared on glass substrates with the microchannel-patterns made of chromium thin films, exposing the SU-8 to UV-light through the pattern. The nominal channel width and depth of the microchannels were 50 μm and 35 μm, respectively. We designed 3 μm-wide and 5 μm-long hydrodynamic traps.

Before each experiment, we preconditioned the microchannel by filling the inlet and outlet wells with washing solutions and applying vacuum at the waste well. Our washing process was as follows: 1 M NaOH for 1 min, 1 M HCl for 1 min, and deionised (DI) water for 1 min. All washing solutions contained 0.1% Triton X-100 to suppress bubble clogging in the hydrophobic microchannel.

Following this, we loaded 9.5 μL of LE and TE to the outlet and inlet wells, respectively, and briefly applied vacuum to the waste well to exchange the solution in the microchannel with LE and TE. The hydrodynamic pressure induced by buffers in the inlet and outlet wells created a pressure driven laminar flow from both inlet and outlet wells toward the waste well and formed a stable LE-TE interface at the junction of three microchannels. We then loaded a 1 μL cell suspension containing a single cell into the inlet well and introduced it into the microchannel via the pressure driven flow. Once we visually confirmed the captured single cell at the hydrodynamic trap (Fig. 1b), we added 9.5 μL of the TE to the waste well to reduce the pressure driven flow. We placed 300 μm diameter platinum wire electrodes into the wells and applied −150 V, −170 V and 0 V to the electrodes at the inlet, waste and outlet wells, respectively. The DC voltage created a concentrated electrical field at the hydrodynamic trap (Supplementary Fig. 1d) and lysed the cytoplasmic membrane within 1 s. Appropriate placement of ITP buffers with the DC electrical field enabled an immediate transition from the lysis to an ITP process that collects and focuses cytoplasmic RNA into an ITP-zone, TE-to-LE interface. At 40 s, we changed the voltages to −350 V and −510 V at the inlet and waste wells, respectively, to accelerate the migration of the ITP-zone. The ITP-zone transported the cytRNA to the output well in about 100 s while the nucleus retained at the hydrodynamic trap. We also monitored current during the extraction with a computer running a custom MATLAB (Mathworks, Inc.) script. The magnitude of the current conducting decreased as the ITP-zone (containing the focused cytRNA) advanced in the channel and as the lower conductivity TE replaced the higher-conductivity LE (Supplementary Fig. 1c). The current signal plateaued near *t* =100 s, coincident with the time at which the focused cytRNA eluted into the outlet well. We deactivated the voltages at 200 s and used a standard pipette to transfer two aliquots from the chip: 9.5 μL from the outlet well containing the cytRNA and 1 μL containing the cell nucleus from the inlet well. Detailed descriptions of a similar protocol and chip, together with a narrated video description, were reported by Kuriyama et al.^15^

### Library preparation and mapping analysis

We synthesised respective cDNA libraries from the fractionated cytRNA and nuclear RNA separately using Smart-seq2 (SMART-seq v4 Ultra Low Input RNA Kit for Sequencing, Clontech) with 18 PCR cycles followed by purification with Agencourt AMPure XP (BeckmanCoulter). We examined the yield and quality of cDNA, respectively, with Qubit 2.0 Fluorometer (ThermoFisher Scientific) and qPCR targeting *GAPDH* (glyceraldehyde-3-phosphate dehydrogenase, Hs02758991_g1, ThermoFisher Scientific) and *HBG* (gamma-globin genes, Hs00361131_g1, ThermoFisher Scientific). We performed the tagmentation reaction with 200 pg cDNA using Nextera XT DNA sample prep kit (Illumina) and purified the cDNA library following the manufacturer’s protocol, except we eluted the cDNA sample with 24 μL instead of 50 μL (see Supplementary Fig. 2a for yields of cDNA). We pooled 98-108 libraries and sequenced these on an Illumina HiSeq2500 with 100-base paired-end reads to an average depth of 4.64 million reads (Supplementary Fig. 2b, c). We mapped the trimmed sequencing reads to the transcripts derived from the human reference genome (GRCh37.75) using a STAR(ver.2.5.1b) of mapping program[31] with ENCODE options, and calculated expression estimates with TPM using RNA-seq by expectation maximisation (RSEM ver.1.3.0)[32]. The average transcriptomic alignments were 94±1% (mean±s.d.) and 93±1%, respectively, with cytRNA-seq and nucRNA-seq (Supplementary Fig. 2d).

### Analysis of intron retention

We computed intron expressions with fragments per kilobase of intron per million mapped reads (FPKM) using 347,041 unique introns (longer than 50 nt) on the genome annotation with 48 SINC-seq data of K562 cells under a standard culturing condition. On average, SINC-seq yielded 14.8% reads mapped to introns with nucRNA-seq, but only 1.1% with cytRNA-seq. Further, SINC-seq detected 37,900±10,800 and 33,800±8,080 unique introns in nucRNA-seq and cytRNA-seq, respectively, and 54,900±9,600 per cell with FPKM of more than 0. We identified an RI that had at least 10% expression level of the gene, 95% coverage in the intronic region, and non-zero expression in the adjacent exon. We discarded intron reads locating on a gene with less than 2 TPM. On the other hand, we identified a fully-spliced intron that had less than 1% expression of the gene and 50% coverage in the adjacent exon. We discarded intron reads that failed the criteria above. We validated the RI identification with splice site scores[33], which showed lower values with RIs than fully-spliced introns (*p*-value < 2.2×10^−16^, U-test), using 9mer (exonic 3mer + intronic 6mer) around 5’ splice site, and 23mer (intronic 20mer + exonic 3mer) around 3’ splice site. In total, we detected 17,335 RIs of which 12,950 and 2,177 RIs had a higher probability of RI in nucRNA and cytRNA, respectively. The RI’s enrichment in the nucleus was consistent with the previous studies[22–24].

To identify the NRI, we calculated the probability of intron retention defined as the proportion of cells with the RI and identified NRI that had 0.25 higher probability in the nuclear fraction than in the cytoplasmic fraction. We then filtered unique NRIs discarding smaller NRIs that had an overlap with a long NRI. On the other hand, we identified CRI that had 0.25 higher probability in the cytoplasmic fraction than in the nuclear fraction.

### Computing cyt-vs nuc-normalised data: in-silico single-cell normalisation

We computed in-silico single-cell RNA-seq data with cytRNA-seq and nucRNA-seq data, scaling the raw TPM values and combining the cyt and nucRNA-seq data as

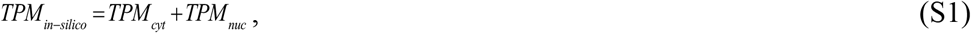

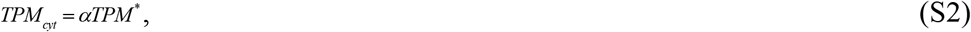

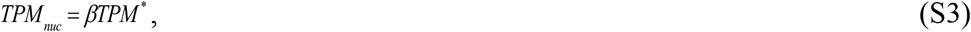

where *α* tand *β* are normalisation factors, which make the summation of the TPM values, *TPM*_*in-siiico*_ of the in-silico single cell data to be one million. We here write raw TPM values with an asterisk and TPM values of an in-silico single cell data, cytRNA-seq, and nucRNA-seq, respectively, with their subscripts. We calculate *α* and *β* using Δ*Ct* of qPCR data taken at the QC of cDNA (Supplementary Data 1) as

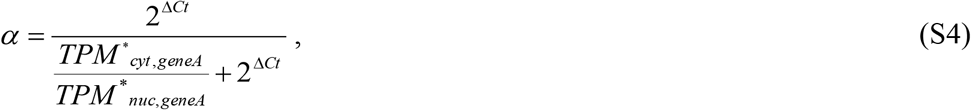

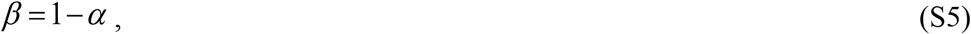

where *TPM*\*_*cyt, geneA*_, *TPM*\*_*nuc, geneA*_, and Δ*Ct* are, respectively, the raw TPM value of *gene A* with cytRNA-seq, the raw TPM value of *gene A* with nucRNA-seq, and Δ*Ct*=*Ct*_nuc, geneA_-*Ct*_cyt, geneA_, which is Δ*Ct* with respect to *gene A*. We calculated pairs of *α* and *β* with *GAPDH* and *HBG* genes (*HBG1*+*HBG2*) as *gene A*s, and used the mean *α* and *β* to compute the *TPM*.

### L-PCA

The PCA for the conventional scRNA-seq that uses a (*n*×*m*) matrix of gene expressions (*n*: number of genes) with multiple samples (*m*: number of samples), however, our L-PCA uses a (2*n*×*m*) matrix of gene expressions, having a dimension by two-fold compared to the conventional, derived from cytRNA-seq and nucRNA-seq. The L-PCA was performed PCA with prcomp in R.

## Declarations

## Acknowledgments

Not applicable

## Authors’ contributions

M.N.A. performed the experiments; Y.O., K.I., H.S., and H.N. analysed the data. S.U., R.Y., and H.K. helped to analyse the data. H.S. and J.G.S. designed and supervised the project. All authors wrote the manuscript.

## Ethics approval

Does not apply for this study.

## Data availability

The data from this study have been deposited into the Sequencing Read Archive and are available for download under accession SRP119800.

---

**Supplementary Figure 1.**
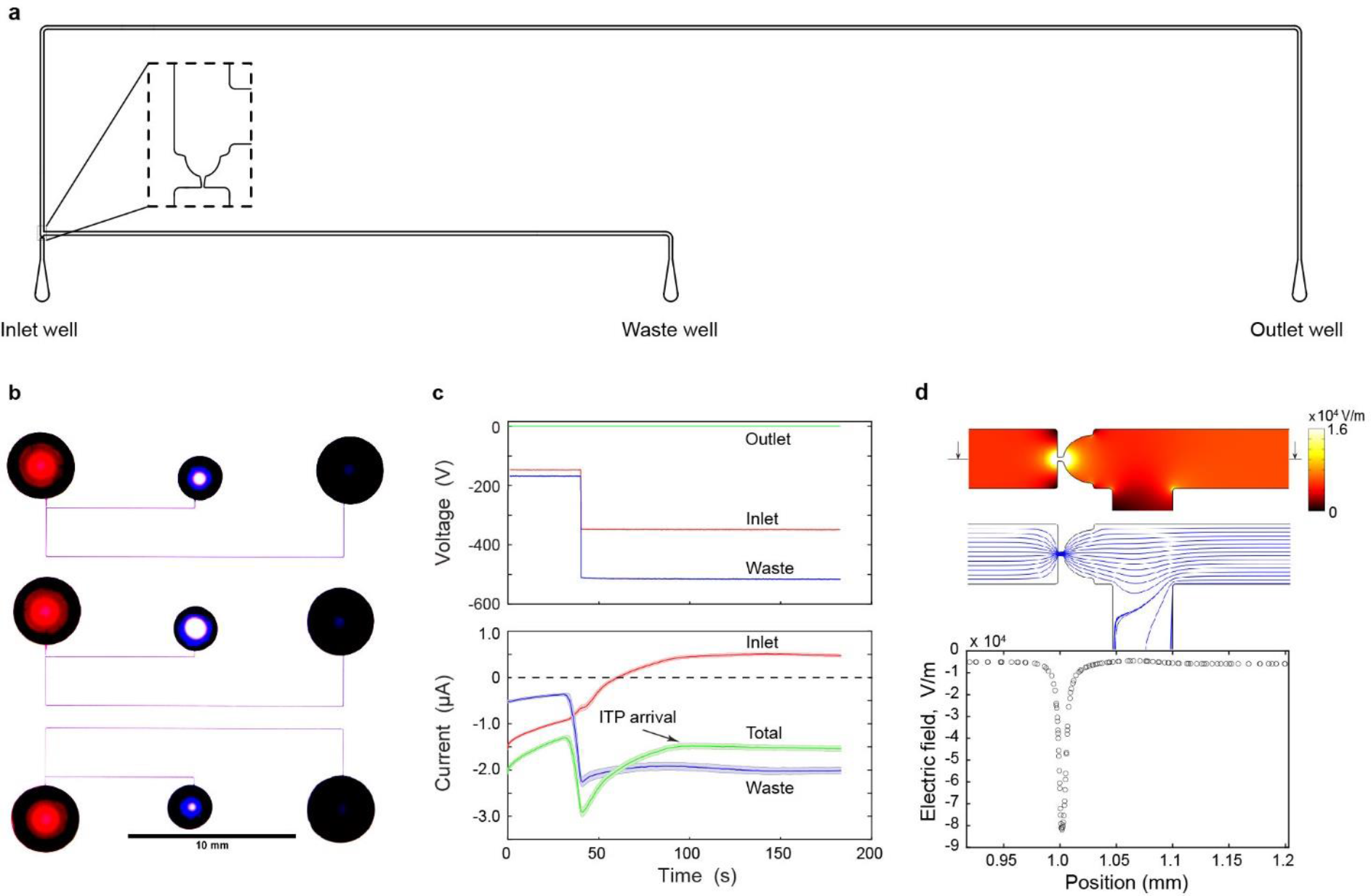
Microfluidic system for SINC-seq protocol. a, Geometry of microfluidic channel with a hydrodynamic trap. b, Photograph of the microfluidic chip consisting of three channels filled with food colouring. c, DC voltage (upper panel) and electrical current behaviour (lower panel) during the extraction protocol. The shade shows 95% confidence interval computed with 48 experiments. d, Magnitude of electrical field around the hydrodynamic trap (top panel) simulated with COMSOL, electrical field lines concentrating at the trapping site (middle panel), and 15-fold electric field amplification at the trap (bottom panel).

**Supplementary Figure 2.**
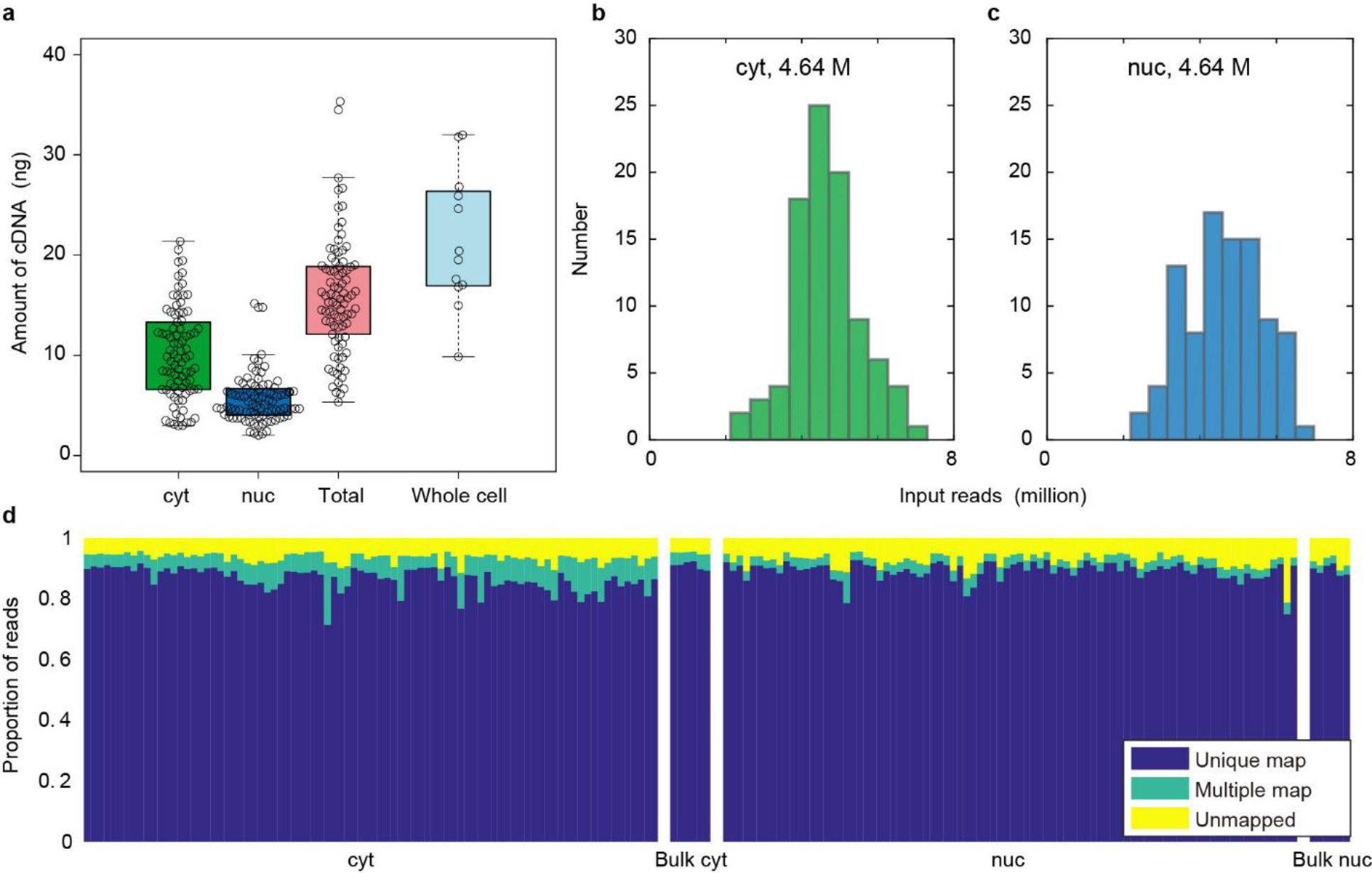
Quality control data of SINC-seq. a, Yields of synthesised cDNA with cytRNA and nucRNA. The total amount of cDNA was calculated by summing cDNA amounts of cytRNA and nucRNA to compare with conventional single-cell protocol. b, c, Input reads of cytRNA-seq and nucRNA-seq. d, Proportion of reads of unmapped, multi-mapped, and uniquely mapped to the GRCh37.75 reference genome.

**Supplementary Figure 3.**
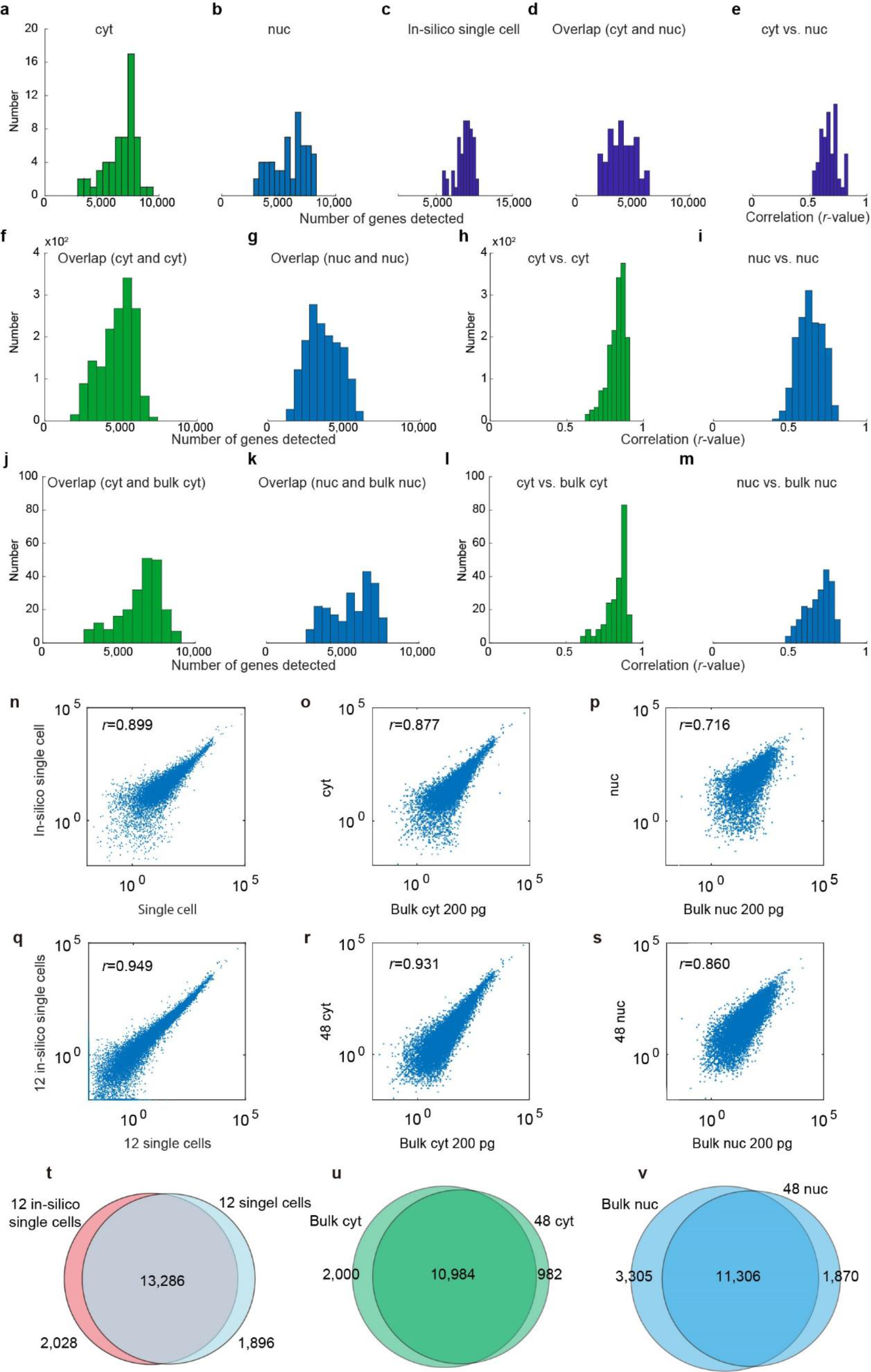
Benchmark of SINC-seq with detection of genes. a-c, Numbers of detected genes with cytRNA, nucRNA and in-silico single cell data, respectively. d, Numbers of genes overlapped in cytRNA and nucRNA. e, Coefficient of correlation between cytRNA and nucRNA. f, g, Numbers of repeatedly detected genes in a pair of cytRNA and in a pair of nucRNA, respectively. h, i, Coefficients of correlation with a pair of cytRNAs and with a pair of nucRNAs. j, k, Numbers of detected genes overlapped between cytRNA-seq and bulk cytRNA-seq, and between nucRNA-seq and bulk nucRNA-seq, respectively. l, m, Coefficients of correlation between cytRNA-seq and bulk cytRNA-seq, and between nucRNA-seq and bulk nucRNA-seq, respectively. n, Correlation of gene expression between in-silico single-cell data and scRNA-seq; o, cytRNA-seq and bulk cytRNA-seq; p, nucRNA-seq and bulk nucRNA-seq. q, Correlation of gene expression between 12 in-silico single-cell data and 12 scRNA-seq; r, 48 cytRNA-seq and bulk cytRNA-seq; s, 48 nucRNA-seq and bulk nucRNA-seq. t, Venn diagrams of detected genes with in-silico normalised data for 12 single cells and conventional 12 scRNA-seq; u, with 48 cytRNA-seq and bulk cyt RNA-seq; v, with 48 nucRNA-seq and bulk nucRNA-seq.

**Supplementary Figure 4.**
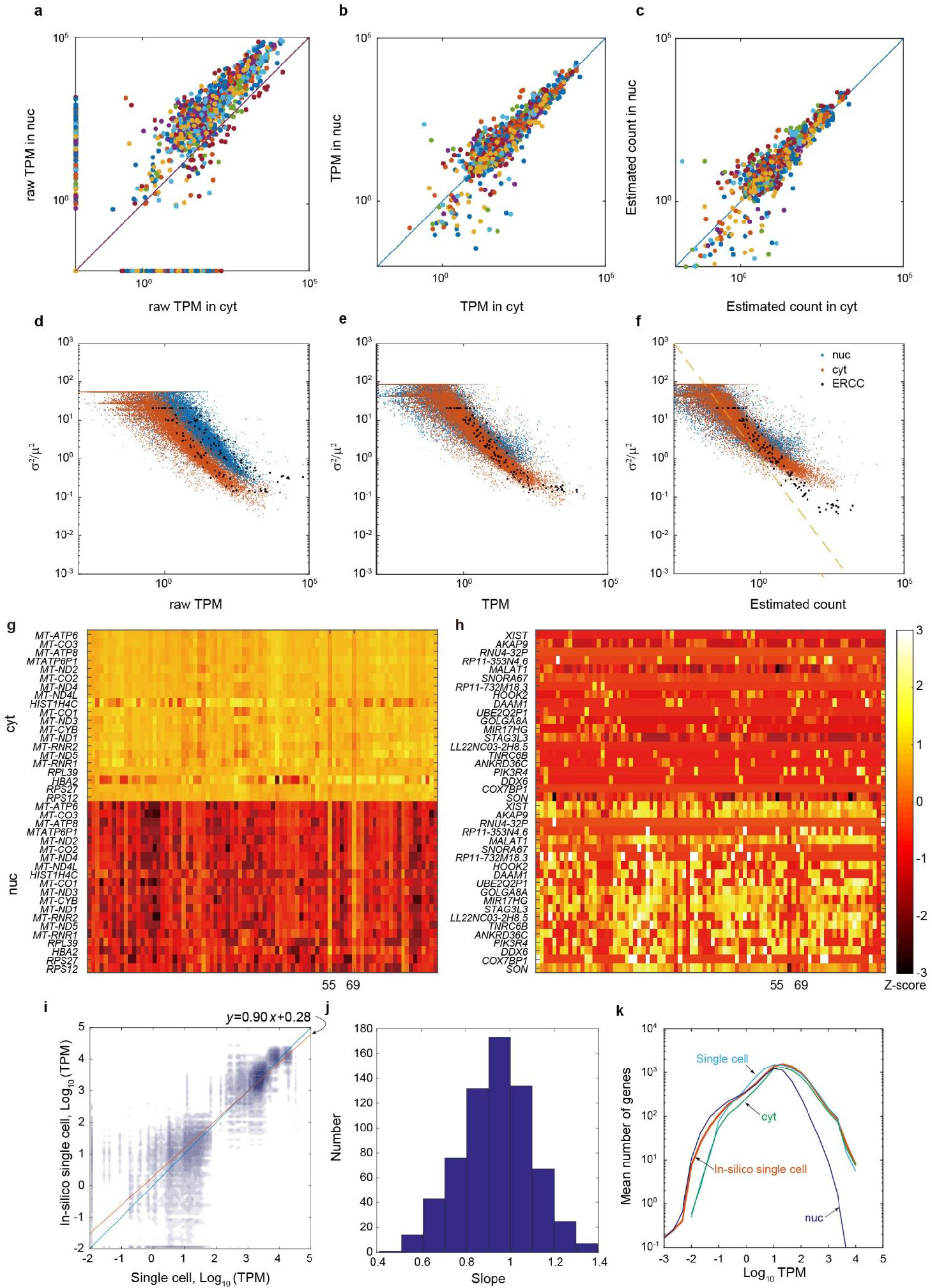
Scaling cytRNA-seq and nucRNA-seq for generating in-silico single-cell data. a-c, ERCC expression comparing cytRNA-seq versus nucRNA-seq with raw TPM, TPM and estimated counts, respectively. d-f, CV^2^ versus mean plots with raw TPM, TPM and estimated counts, respectively. g, h, Fractionation stringency assessed with top 20 genes enriched in cytRNA and nucRNA, respectively. i, Comparison of expression patterns of the top 20 genes enriched in cytRNA and nucRNA between in-silico single-cell data and scRNA-seq. j, Statistics of the slope comparing in-silico single cell data versus scRNA-seq with the expression level of the top 20 enriched genes. k, In-silico single cell data shows a wider dynamic range of detected genes integrating cytRNA-seq and nucRNA-seq.

**Supplementary Figure 5.**
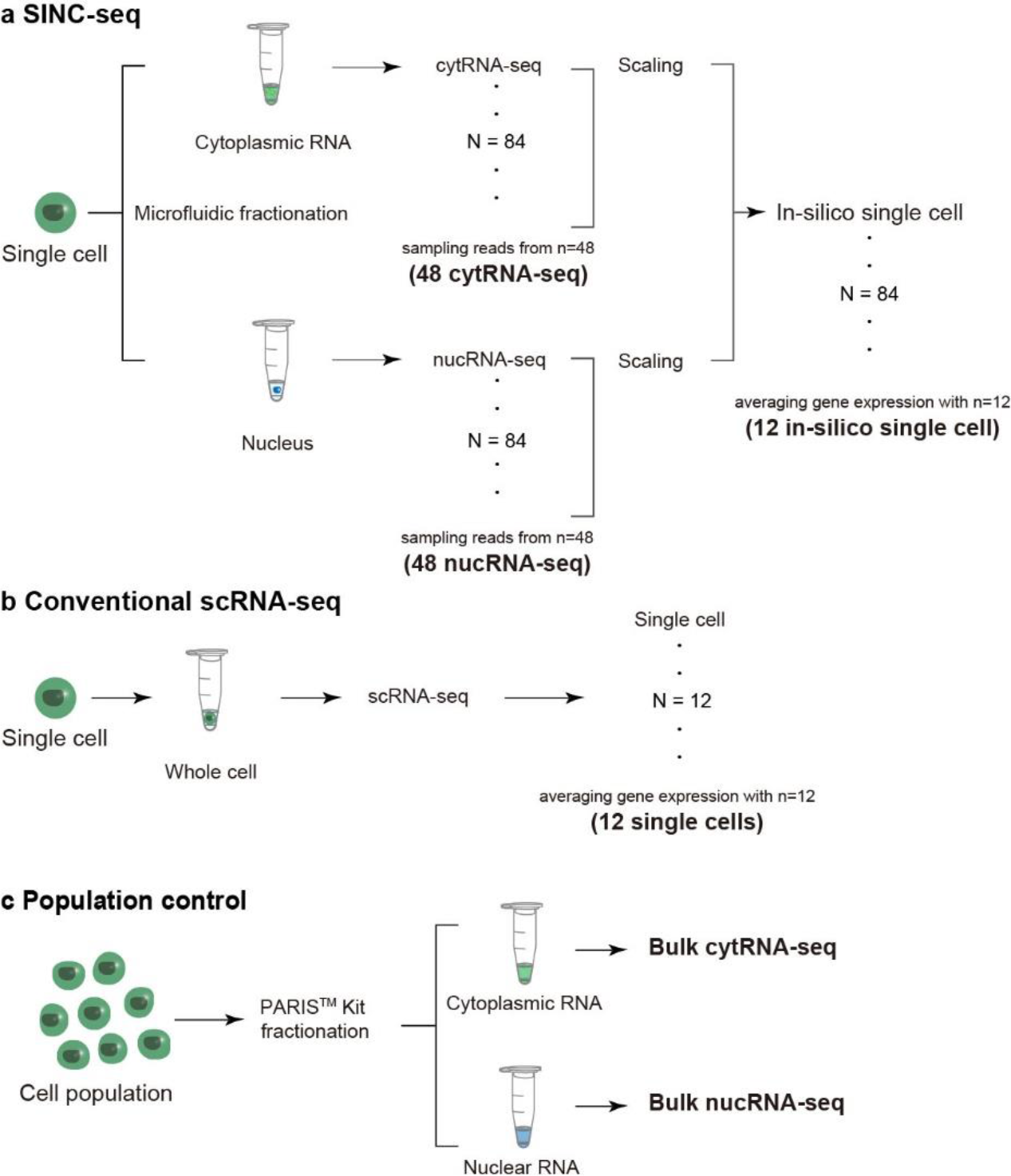
Overview of RNA-seq samples. a, SINC-seq constructs cytRNA-seq and nucRNA-seq per cell with cytoplasmic RNA and a nucleus, respectively. 48 cytRNA-seq and 48 nucRNA-seq data were, respectively, created by randomly sampling reads from 48 of cytRNA-seq and 48 of nucRNA-seq. In-silico single cell data were created by scaling and integrating cytRNA-seq and nucRNA-seq from the same single cell. The 12 in-silico single cell data was created by averaging 12 of randomly sampled in-silico single cell data sets. b, Conventional scRNA-seq. 12 single cell RNA-seq was created with averaging the 12 of scRNA-seq. c, Population controls of bulk cytRNA-seq and bulk nucRNA-seq were prepared with PARIS Kit, which fractionates cytoplasmic RNA and nuclear RNA with a population of cells, followed by Smart-seq2 protocol with 200 pg RNA and 15 PCR cycles.

**Supplementary Figure 6.**
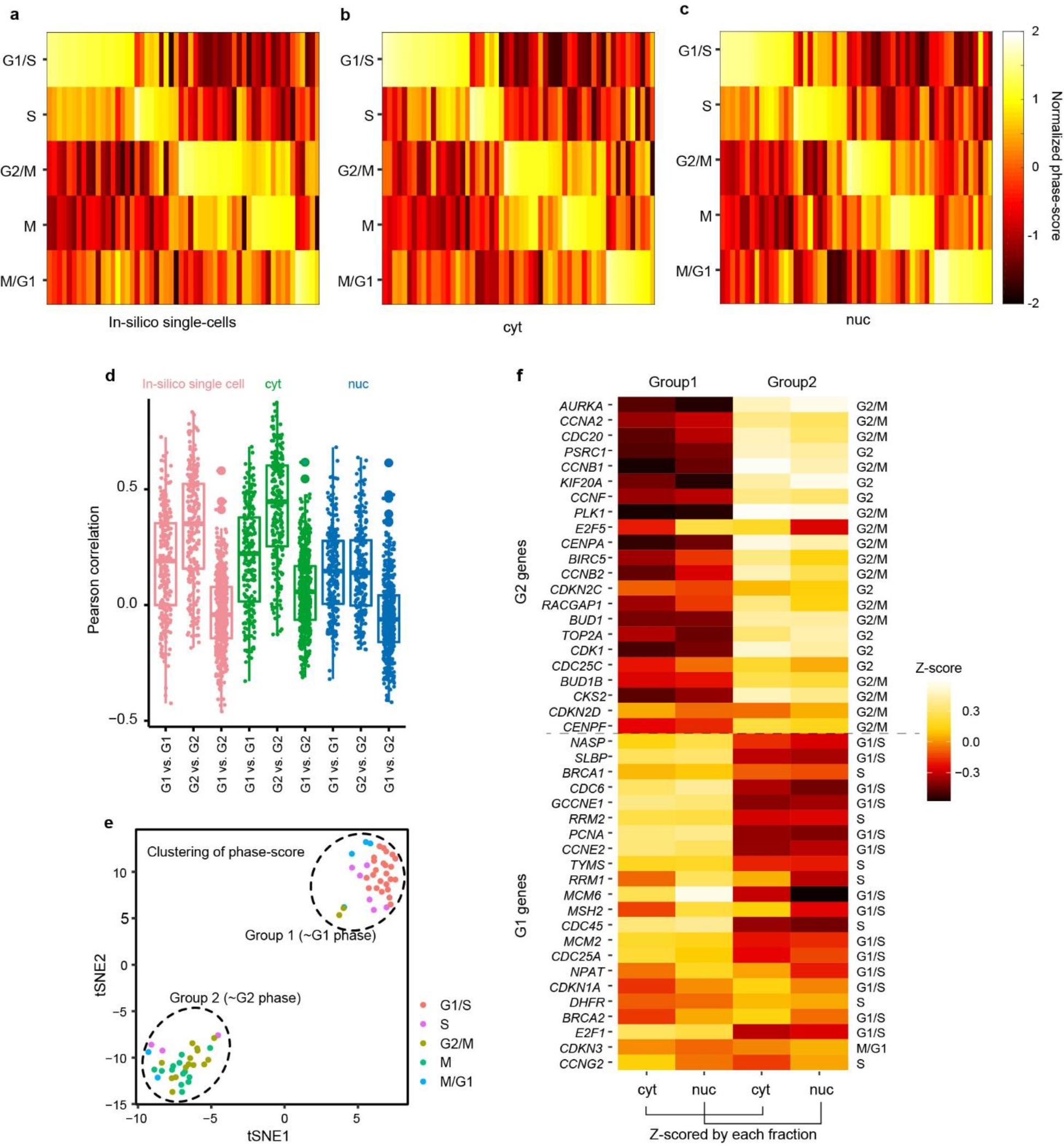
Cell-cycle phase of K562 cells analysed by SINC-seq. a-c, Normalized phase scores of in-silico single cell data, cytRNA-seq, and nucRNA-seq, respectively. d, Pearson correlations shown in Fig. 2 d-f. e, tSNE with normalised phase scores (Supplementary Fig. 6a) segregates cells into G1 and G2 groups. f, Z-score calculated with individual fraction indicates relative expression of each gene among G1 and G2 groups.

**Supplementary Figure 7.**
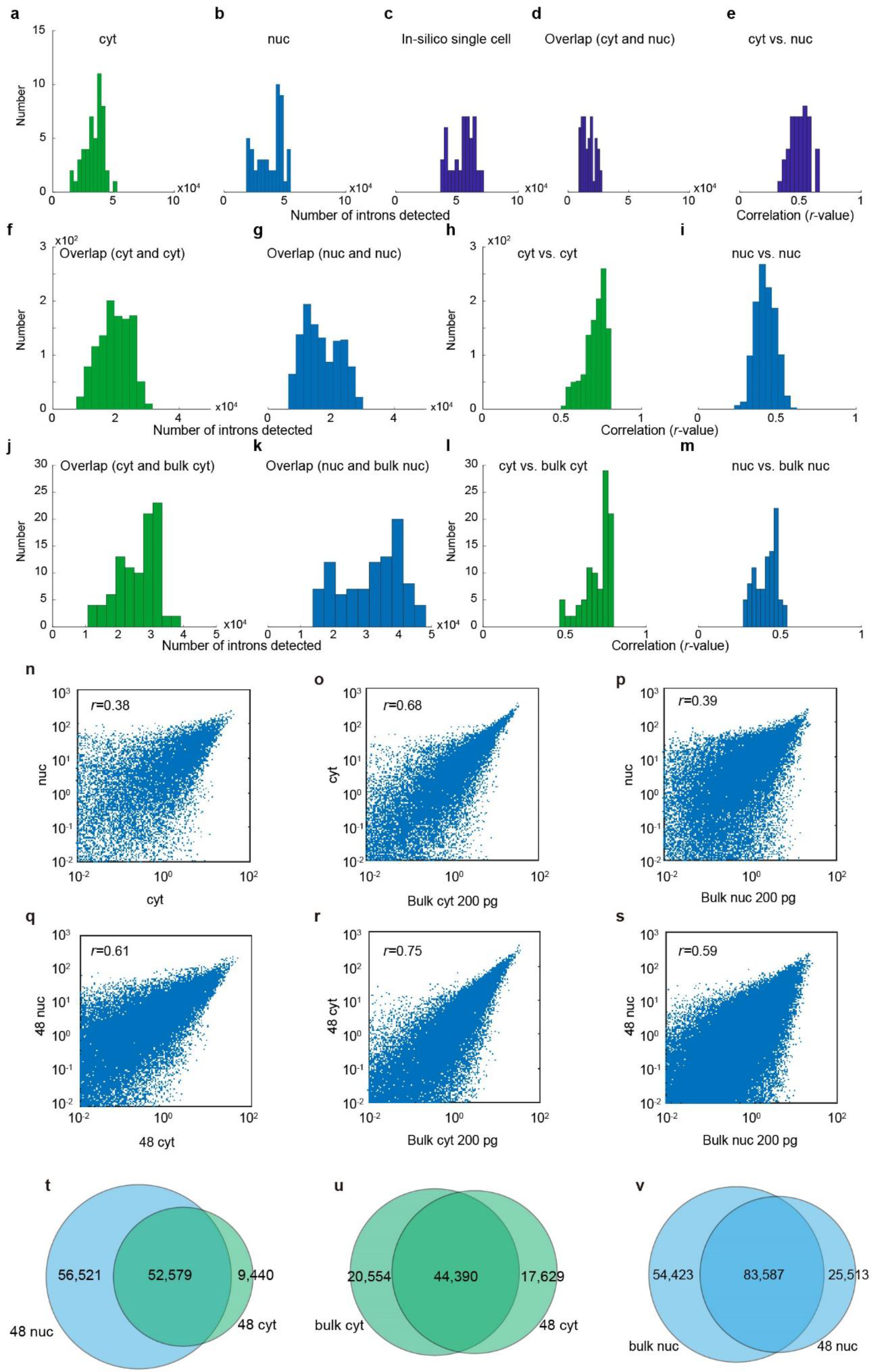
Benchmark of SINC-seq with detection of introns. a-c, Numbers of detected introns with cytRNA, nucRNA and in-silico single cell data, respectively. d, Number of detected introns overlapped in cytRNA and nucRNA. e, Coefficient of correlation with intron abundance between cytRNA and nucRNA. f, g, Numbers of detected introns in a pair of cytRNAs and in a pair of nucRNAs, respectively. h, i, Coefficients of correlation with a pair of cytRNAs and a pair of nucRNAs, respectively. j, k, Numbers of detected introns overlapped cytRNA-seq and bulk cytRNA-seq, and nucRNA-seq and bulk nucRNA-seq, respectively. l, m, Coefficients of correlation between cytRNA-seq and bulk cytRNA-seq, and between nucRNA-seq and bulk nucRNA-seq, respectively. n, Correlation of intron abundance between cytRNA-seq and nucRNA-seq; o, cytRNA-seq and bulk cytRNA-seq; p, nucRNA-seq and bulk nucRNA-seq. q, Correlation of intron abundance between 48 cytRNA-seq and 48 nucRNA-seq; r, 48 cytRNA-seq and bulk cytRNA-seq; s, 48 nucRNA-seq and bulk nucRNA-seq. t, Venn diagrams of detected introns with 12 cytRNA-seq and 12 nucRNA-seq; u, with 48 cytRNA-seq and bulk cytRNA-seq; v, with 48 nucRNA-seq and bulk nucRNA-seq.

**Supplementary Figure 8.**
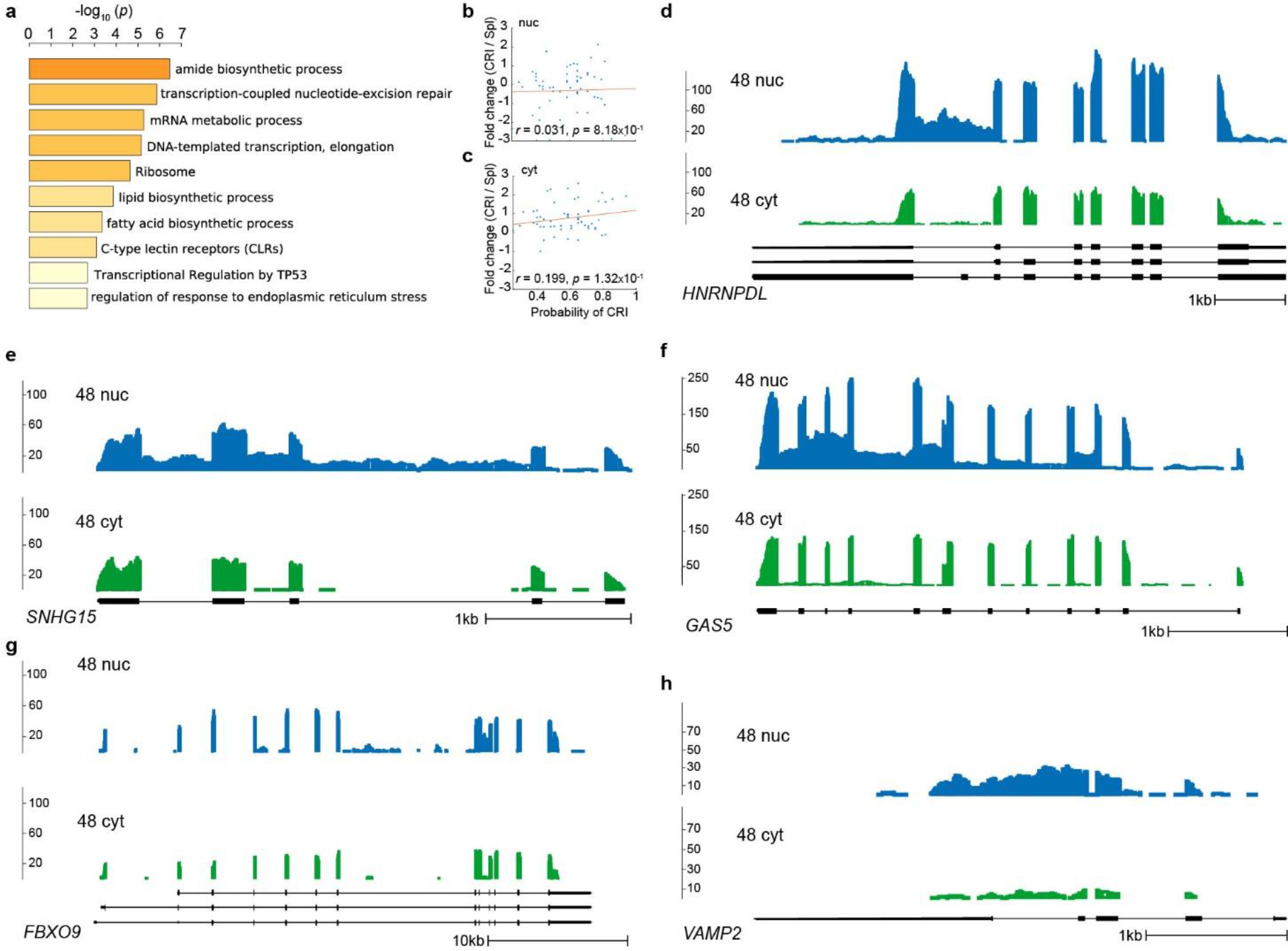
NRI and CRI have different enriched functions. a, Gene ontology analysis with CRI. b, c, Correlation analysis between the probability of CRI and the fold change of gene expression among cells with CRI and without CRI (Spl: spliced) in nucRNA and cytRNA, respectively. Coverages of d, *HNRNPDL* e, *SNHG15* f, *GAS5* (*SNHG2*), g, *FBXO9*, and h, *VAMP2* genes.

**Supplementary Figure 9.**
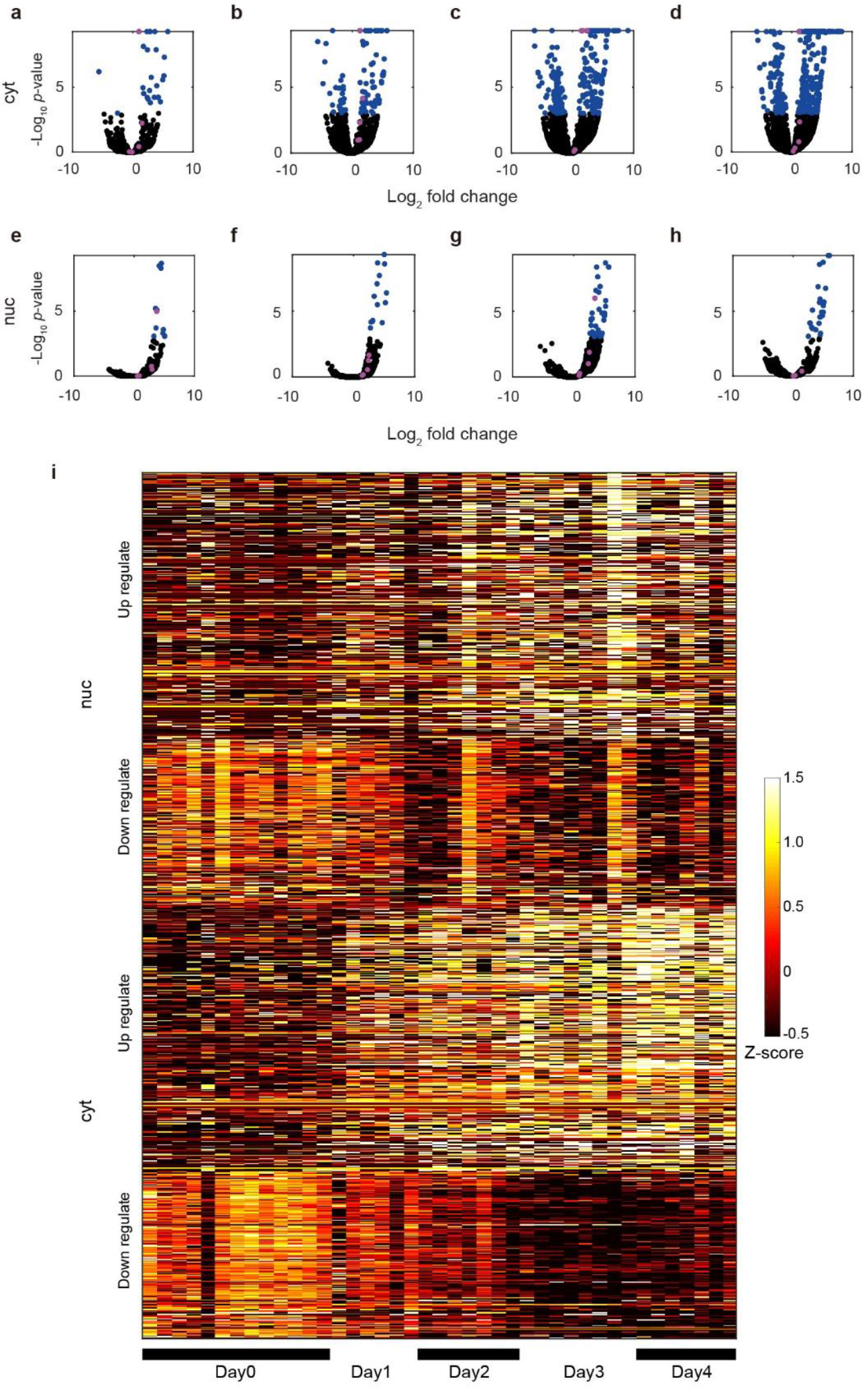
Differentially expressed genes along sodium butyrate-induced K562 differentiation. a-d, Differential expression with cytRNA of day 0 vs. day 1, day 0 vs. day 2, day 0 vs. day 3, and day 0 vs. day 4, respectively. e-h, Differential expression with nucRNA of day 0 vs. day 1, day 0 vs. day 2, day 0 vs. day 3, and day 0 vs. day4, respectively. We identified DEG using “nbintest” and “mafdr” of MATLAB functions. Blue, genes with *p* values less than 0.001 and absolute log2 fold changes greater than unity. Pink, *GATA1*, *HBG1*, *HBG2*, *GYPA* and *TFRC* genes. i, Heatmap showing up- and down-regulations of DEG in cytRNA and nucRNA.

**Supplementary Figure 10.**
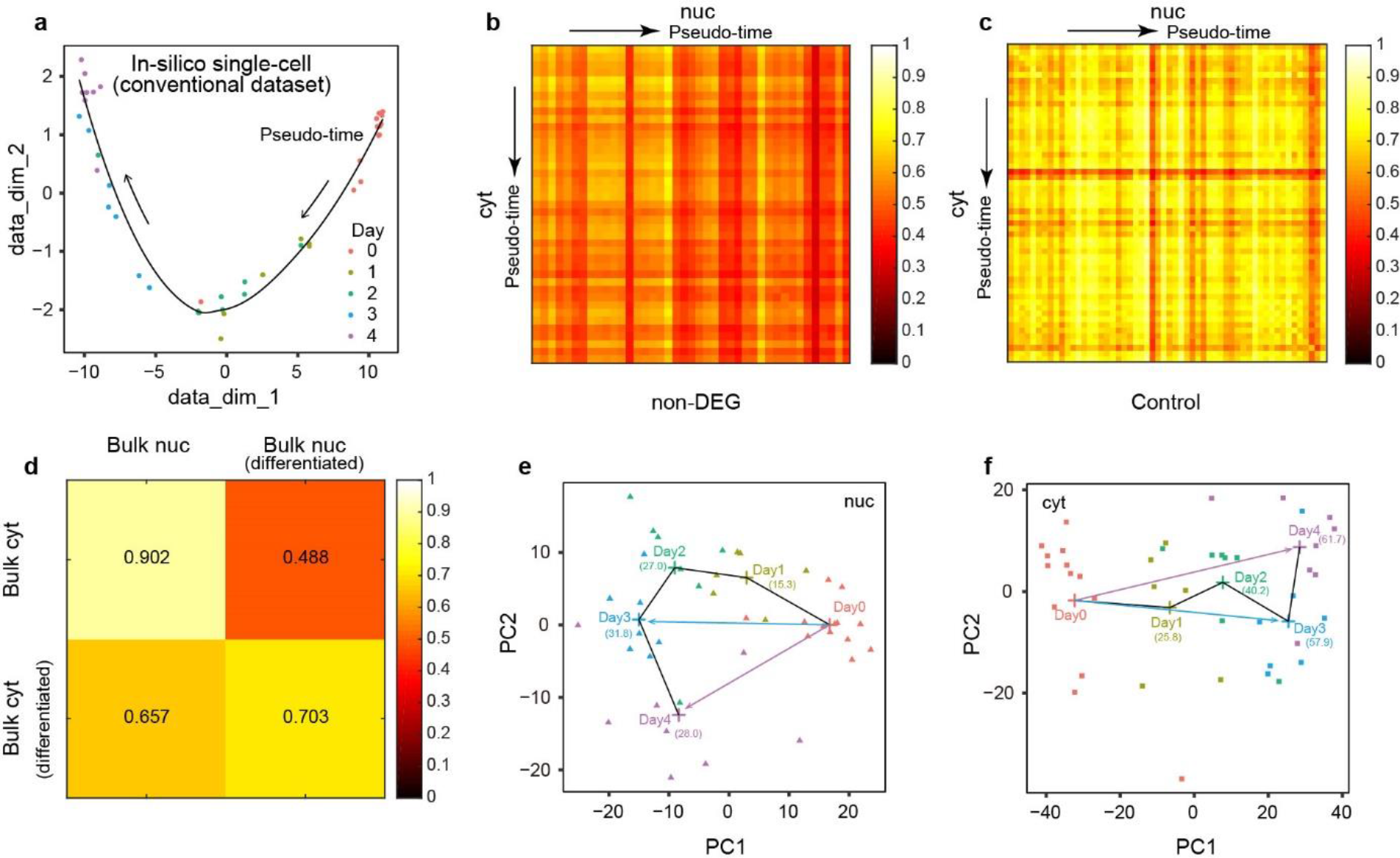
Correlation dynamics under sodium butyrate differentiation of K562 cells. a, Pseudo-time computed with Monocle2^27^ using DEG expression of in-silico single-cell data. b, Cross-correlation between cytRNA and nucRNA computed with non-DEG of differentiating K562 cells. c, Cross-correlation of cytRNA and nucRNA computed with DEG of non-differentiating K562 cells along pseudo-time. d, Cross-correlation of bulk cytRNA-seq and bulk nucRNA-seq computed with DEG of differentiating K562 cells. e, f, PCA on cytRNA-seq and nucRNA-seq data of differentiating K562 cells, respectively. Cross-points in these panels indicate the centre of mass of each cluster.

